# Precision genome-editing with CRISPR/Cas9 in human induced pluripotent stem cells

**DOI:** 10.1101/187377

**Authors:** John P. Budde, Rita Martinez, Simon Hsu, Natalie Wen, Jason A. Chen, Giovanni Coppola, Alison M. Goate, Carlos Cruchaga, Celeste M. Karch

**Author notes:** authors contributed equally to this work (John P. Budde, Rita Martinez). Correspondence/Lead Contact: Celeste M Karch, Department of Psychiatry, Washington University School of Medicine, 425 S. Euclid Ave, Campus Box 8134, St. Louis, MO 63110.

## Abstract

Genome engineering in human induced pluripotent stem cells (iPSCs) represent an opportunity to examine the contribution of pathogenic and disease modifying alleles to molecular and cellular phenotypes. However, the practical application of genome-editing approaches in human iPSCs has been challenging. We have developed a precise and efficient genome-editing platform that relies on allele-specific guideRNAs (gRNAs) paired with a robust method for culturing and screening the modified iPSC clones. By applying an allele-specific gRNA design strategy, we have demonstrated greatly improved editing efficiency without the introduction of additional modifications of unknown consequence in the genome. Using this approach, we have modified nine independent iPSC lines at five loci associated with neurodegeneration. This genome-editing platform allows for efficient and precise production of isogenic cell lines for disease modeling. Because the impact of CRISPR/Cas9 on off-target sites remains poorly understood, we went on to perform thorough off-target profiling by comparing the mutational burden in edited iPSC lines using whole genome sequencing. The bioinformatically predicted off-target sites were unmodified in all edited iPSC lines. We also found that the numbers of *de novo* genetic variants detected in the edited and unedited iPSC lines were similar. Thus, our CRISPR/Cas9 strategy does not specifically increase the mutational burden. Furthermore, our analyses of the *de novo* genetic variants that occur during iPSC culture and genome-editing indicate an enrichment of *de novo* variants at sites identified in dbSNP. Taken together, we propose that this enrichment represents regions of the genome more susceptible to mutation. Herein, we present an efficient and precise method for allele-specific genome-editing in iPSC and an analyses pipeline to distinguish off-target events from *de novo* mutations occurring with culture.

## Introduction

For many neurodegenerative diseases, rare mutations lead to autosomal dominant forms of the disease. Increasingly, large genetic studies of sporadic cases of neurodegenerative disease reveal novel variants that modify disease risk. Thus, establishing cellular models that correct or introduce these pathogenic mutations and risk variants is critical for understanding the molecular mechanisms that underlie disease.

Clustered Regularly Interspaced Short Palindromic Repeats (CRIPSR)/Cas has emerged as the most versatile method for modifying the genome in mice, cells, and other model organisms. CRISPR/Cas has two components: (1) Cas9 endonuclease and (2) guide RNA (gRNA) (1). gRNAs direct the Cas9 endonuclease to specific sites in the genome where single-or double-stranded breaks trigger natural DNA repair processes such as non-homologous end joining (NHEJ) or homologous recombination (HR) in the presence of a donor DNA template (2). NHEJ events, which typically produce insertions/deletions (INDELs), occur frequently and have been exploited to generate gene knockouts. HR is necessary for single allele editing; yet, HR events occur at low frequencies. As such, the specificity and efficiency of the gRNA to direct HR remains a limitation of effectively implementing precision editing of a single allele (3-6).

Limitations in controlling gRNA specificity and Cas9 activity have made precision genome-editing of single alleles in human iPSC largely inefficient. Furthermore, inherent variability in individual iPSC donor lines and their ability to maintain pluripotency throughout the editing pipeline limits the broader application of this approach. Several methods have been reported to improve editing efficiency and specificity by modifying the Cas9 and gRNA structure (6-12) or isolation and analysis of edited clones (13,14). Yet, these improvements alone have not been sufficient to enable the broader use of precision editing of a single allele in iPSCs.

Efficient precision genome-editing relies on highly specific gRNAs paired with a robust method for culturing and screening the modified iPSC clones. We have developed an efficient precision genome-editing platform. By applying an allele-specific gRNA design strategy, we have demonstrated greatly improved editing efficiencies without the introduction of additional modifications of unknown consequence in the genome (e.g. antibiotic selection or synonymous changes in donor sequences), making our approach seamless. Applying this allele-specific gRNA design alongside a strictly defined pipeline for nucleofection and clone screening, we are able to obtain iPSC modified at a single allele within one month. We have applied this approach to generate isogenic clones from nine independent donor iPSC at five independent loci across multiple chromosomes.

With an efficient method for allele-specific editing in hand, we sought to more carefully assess precision by investigating off-target effects of CRISPR/Cas9, which remain poorly understood. While careful gRNA design focused on minimizing sequence homology at other sites in the genome can reduce off-target changes in the genome (15), recent evidence suggests that factors beyond sequence homology affect which sites are changed (12,16). Furthermore, recent studies demonstrate that selective pressures induced during passage are sufficient to induce *de novo* changes in p53 and other cancer genes in iPSCs (17-19). Thus, we sought to apply careful downstream analysis to determine the extent to which CRISPR/Cas9 alters the genome-wide mutational burden on edited iPSCs. To evaluate genome-wide mutational burden, we generated whole genome sequencing (WGS) data for iPSCs that underwent genome-editing, parental iPSC and parental fibroblasts. We did not observe enrichment in off-target changes in the edited lines. We found that the mutational burden within the genome-edited iPSC cultures is likely due to well-established selective pressures that occur with culture rather than CRISPR/Cas9 induced off-target effects. Among the variants identified, we observed enrichment in *de novo* variants in sites that are previously reported in dbSNP. We propose that these variants represent regions of the genome that are most susceptible to change.

## Results

### Novel pipeline for allele-specific genome-editing in iPSC

In order to define the molecular and cellular consequences of variants that underlie risk for neurodegenerative disease, we sought to develop an efficient and precise approach for editing disease associated alleles in human iPSCs. Single allele editing relies on HR events, which occur at low frequency. Thus, we focused on adaptation of gRNA design and iPSC culturing to increase the introduction of the desired modification (Figure 1).

**Figure 1:**
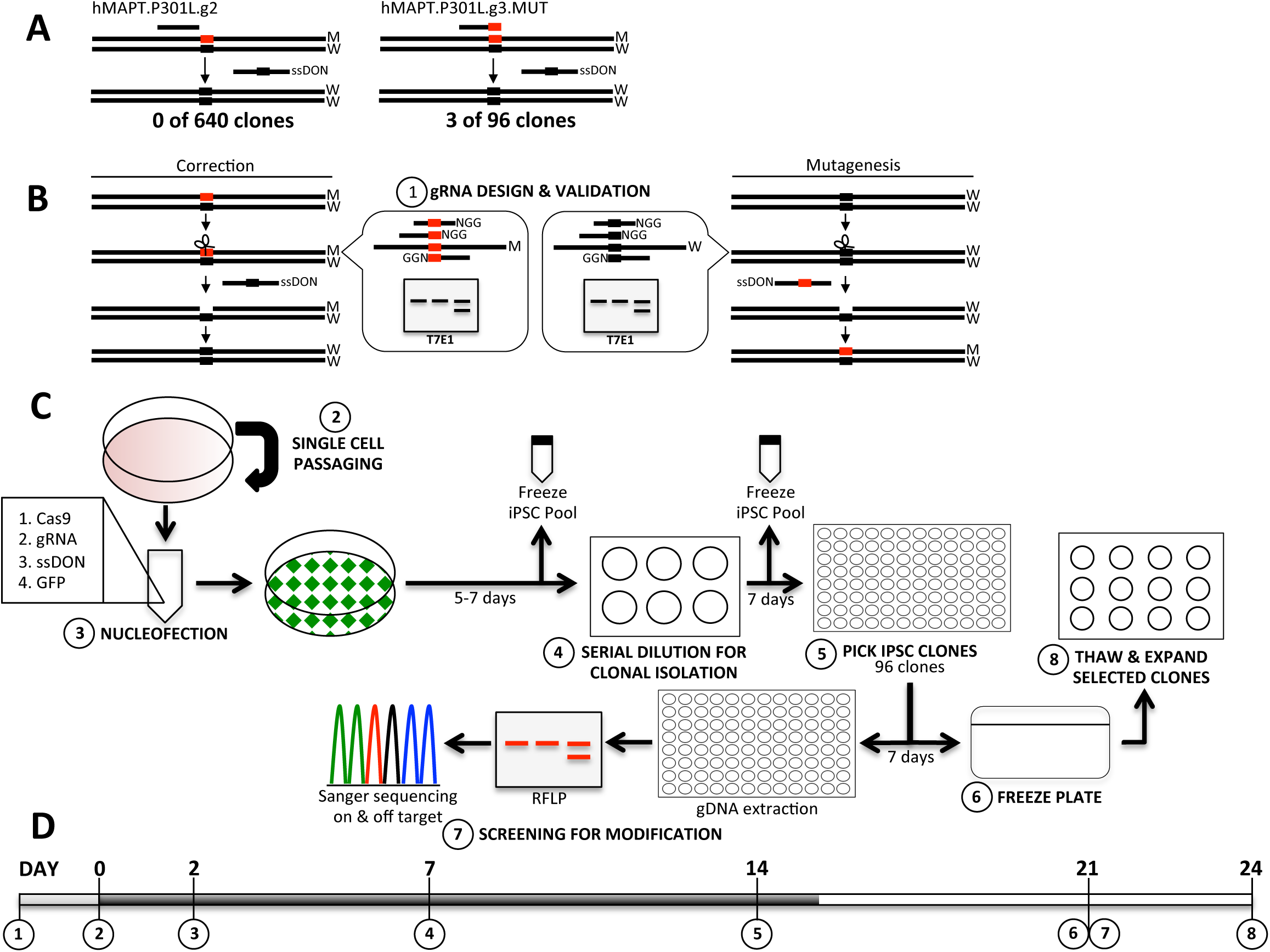
Workflow for efficient genome-editing in human iPSC. A. Comparison of the design strategy and resulting efficiencies for allele-specific and non-specific gRNA. B. Allele-specific gRNA design for correction (left) and mutagenesis (right). Inset demonstrates location of allele to be modified within gRNA sequence. C. Work flow for genome-editing. D. Timeline for editing. Numbers represent major steps in the editing pipeline. M, mutant allele. W, wild-type allele.

Our genome-editing efforts were focused on correcting patient iPSC lines that are heterozygous for dominantly inherited mutations that cause frontotemporal dementia. We applied a standard pipeline for designing gRNAs (15). gRNAs were prioritized based on identifying PAM sequences within 20bp of the target locus and minimizing sequence homology in the genome (minimal off-targets based on 1, 2, or 3 mismatches in the genome). Using this approach, we designed a gRNA 1bp upstream of the target allele with minimal off-targets (Table 1; Figure 1A; hMAPT.P301L.g2). This gRNA produced high NHEJ frequency (23%) in a T7E1 mismatch assay in K562 cells, demonstrating the gRNA is active. After nucleofection of Cas9, gRNA, and ssODN in the iPSC (see Methods), screening of more than 600 iPSC clones revealed 100 iPSC clones with NHEJ events (e.g. INDELs) at the target site and 10 clones with modification of the mutant allele in addition to an INDEL at the target site, suggestive of a secondary NHEJ event after HR (Table 1). Thus, this gRNA failed to produce iPSC clones in which the mutant allele was corrected.

**Table 1:** Mutant specific gRNA enhances desired modifications of human iPSC

| Guide name | Guide Length | Target Allele | Mutant Allele | Off Targets° | NHEJ Activity^ | ssODN* | Clones Screened | Clones with Desired Modification |
| --- | --- | --- | --- | --- | --- | --- | --- | --- |
| hMAPT.P301L.g2 | 20 | T | C | 5 | 23% | WT | 640 | 0 |
| hMAPT.P301L.g3.WT | 20 | C | T | 3 | 24% | P301L | 90 | 1 (4<sup>#</sup>) |
| hMAPT.P301L.g3.MUT | 20 | T | C | 3 | 24% | WT | 96 | 3 |
|  |  |  |  |  |  | P301S | 90 | 16 (5<sup>#</sup>) |
| hMAPT.P301L.g3.17.MUT | 17 | T | C | 3 | 4% | WT | 96 | 1 |
| hMAPT.R406W.g11.MUT | 20 | T | C | 38 | 23% | WT | 60 | 2 |
|  |  |  |  |  |  |  | 96 | 1 |
|  |  |  |  |  |  |  | 71 | 1 |
| hMAPT.V337M.g4.MUT | 20 | A | G | 60 | 56% | WT | 96 | 1 |
|  |  |  |  |  |  |  | 86 | 4 |
|  |  |  |  |  |  |  | 76 | 5 |
| hPLD3.A442A.g9.MUT | 20 | A | G | 5 | 20% | WT | 88 | 1 |

^K652 cells. <sup>#</sup>Homozygous for mutation. °Bioinformatically predicted off-target effects based on matches in the genome for short arm of gRNA only. Matches or 1, 2 mismatches in the long arm of gRNA were predicted to exist. \* ssODN, single stranded oligodeoxynucleotide.

In order to increase editing success, we needed to either enhance HR or reduce secondary NHEJ events after HR occurs. We chose to apply an approach aimed at reducing secondary NHEJ events after HR. We designed a gRNA for the same target allele in which the gRNA sequence overlaps with the target allele (Table 1; Figure 1A; hMAPT.P301L.g3.mut). We hypothesized that once HR occurs, the allele-specific gRNA would have lower affinity for the new allele. This gRNA was also active in K562 cells (24%NHEJ frequency). Screening of 96 clones produced three correctly modified iPSC clones (Table 1; hMAPT.P301L.g3.mut). Extending this approach to an additional four sites and in nine independent donor iPSC lines resulted in consistently successful editing (Table 1; Figure 1).

To determine whether we could enhance our editing efficiency further, we examined the impact of truncated gRNAs on single allele editing. Truncation of gRNAs has been reported to enhance gRNA specificity (7,20). Guides truncated to 17 bases of complementary sequence (17mer) have been reported to produce comparable on-target activity to guides with 20 bases of complementary sequence (20mer) (7). Yet, 17mer gRNAs reportedly produce significantly fewer off-target events in immortalized cell lines (7). To test whether allele-specific gRNA design and gRNA truncation would have an additive effect on enhancing specificity and overall efficiency of editing in human iPSCs, we compared allele-specific gRNAs of varying lengths. A truncated gRNA (hMAPT.P301L.g3.17.mut; 17mer) produced lower NHEJ frequency in a T7E1 assay than the full-length gRNA (hMAPT.P301L.g3.mut; 20mer; 4% v. 24%, respectively; Table 1). The truncated gRNA also produced fewer correctly modified iPSC clones than the full-length gRNA (Table 1; hMAPT.P301L.17.g3.mut). Additionally, WGS data demonstrated that the mutational burden in the iPSCs corrected with the 17mer versus 20mer gRNAs was similar (Supplemental Figure 1). Thus, truncated gRNAs at this locus do not enhance allele-specific targeting and editing efficiency.

We next sought to assess the off-target effects of CRISPR/Cas9 in the correctly modified clones (e.g.: *MAPT* P301L to WT). We identified three sites in the genome that match with the short arm of the guide in chr6:10095527, chr12:8545565, and chr21:42619506 by bioinformatic analysis (15) (Table 1). We examined the sequence in the 300bp surrounding these off-target sites. The sequence surrounding each of these sites was identical in all lines analyzed by WGS (Supplemental Figure 2). Our findings from the WGS data were consistent with Sanger sequencing of these predicted off-target sites (Supplemental Figure 2).

### Assessing genome-wide mutational burden in genome edited iPSC

CRISPR/Cas9 may induce changes in the genome beyond the on-target site. Off-target sites can be predicted computationally by comparing sequence similarity of the gRNA to the entire genome (15). However, factors beyond sequence homology also contribute to off-target effects (12). To evaluate mutational burden in CRISPR/Cas9-edited iPSC lines, we performed WGS in 13 cell lines (Figure 2): parental fibroblasts, parental iPSC, unedited iPSC exposed to the editing pipeline, and edited iPSC. WGS data from all iPSC clones was aligned and cleaned. To evaluate *de novo* variants, all comparisons were made with the cell line from the previous generation.

**Figure 2:**
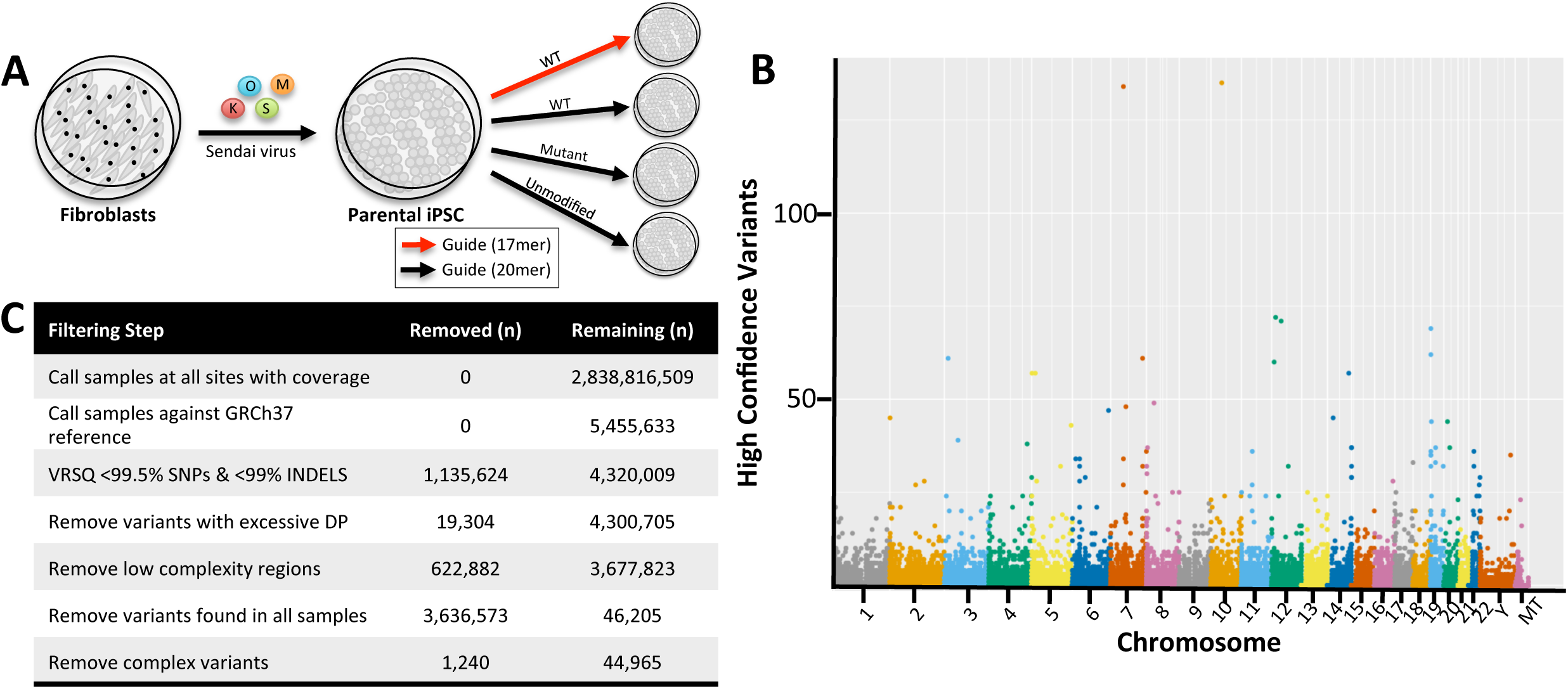
Experimental and computational approach for identifying genomic variation induced by genome-editing. A. Diagram of the derivation of the samples included in WGS. O, Oct4; M, cMyc; K, Klf4, S, Sox2. B. Manhattan plot demonstrating distribution of variants across the genome. C. Diagram detailing the filtering strategy. Next to each filtering step: variant removed/variants remaining.

To assess mutational burden, we applied a pipeline for calling and cleaning the data that was modified from previous studies (17,21). WGS data was aligned to the GRCh37 reference genome. The aligned WGS data was cleaned in a sequential multistep process, guided by the GATK Best Practices (22). After application of VQSR to remove low quality variants, sites with excessive coverage and low-complexity regions of the genome were excluded, as were any genotype calls with depth < 8 or phred-scaled genotype quality < 20 (Figure 2). In previous studies reporting WGS in iPSC clones, the total number of high confidence variants varies greatly (6,17,23-25). This variability is, in part, due to the extent to which the WGS data is interrogated (e.g. analysis of only predicted off-target regions). This variability is also driven by use of an analysis pipeline that includes removal of all variants reported in dbSNP during cleaning. dbSNP is a database of reported genetic variation in humans and other organisms (26). While many entries in dbSNP may be benign, dbSNP contains rare Mendelian mutations, non-synonymous variants, and common variants that increase risk for complex diseases. We propose that when screening for off-target effects, variants reported in dbSNP are as critical to evaluate as any other variant. Thus, our pipeline retains dbSNPs, resulting in a higher total number of variants.

To assess the genome-wide impact of reprogramming on mutational burden, we investigated *de novo* variants in parental iPSC prior to genome-editing (reference: parental fibroblasts; Figure 3). We identified 1,417 *de novo* single nucleotide polymorphisms (SNPs) in the iPSC (reference: fibroblast; Figure 3). If we ignore variants reported in dbSNP as previous pipelines suggest, we identified 690 *de novo*, novel variants. This is lower than previous studies reporting between 1000 and 2000 *de novo*, novel variants in reprogrammed iPSCs (17,18). *De novo* INDEL findings were consistent with single variant findings (Figure 3). Thus, the reprogramming event and subsequent passage of the iPSC culture is sufficient to induce *de novo* changes.

**Figure 3:**
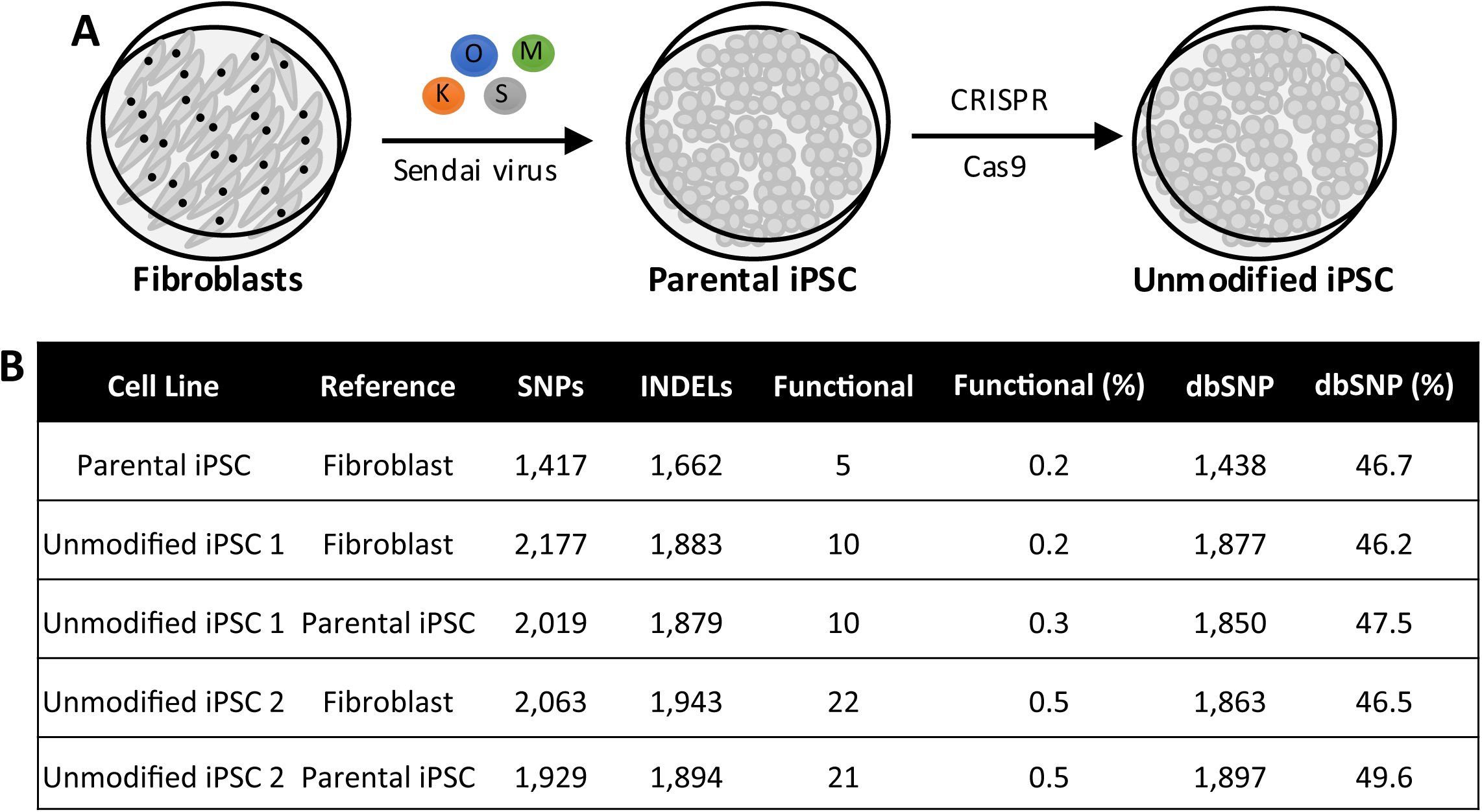
*De novo* variants occurring during transformation of culture. A. Diagram. *De novo* variant counts for cell lines after reprogramming and after exposure to genome-editing pipeline but where no editing occurs. B. Table of counts. Functional variants are defined as synonymous, frameshift, or missense variants in coding regions.

We next sought to more carefully evaluate mutational burden induced by CRISPR/Cas9 beyond the predicted off-target sites, we compared the number of *de novo* variants in the edited iPSC (reference: parental iPSC; Figure 4A; Table 2). Overall, we found that the relative number of *de novo* SNPs were similar across the edited lines, with the exception of M.6 (Figure 4B; Supplemental Table 1). Our findings were similar for *de novo* INDELs. To better understand the types of *de novo* changes, we evaluated the context of the mutations and transversion rates in our samples. We found no appreciable differences among the edited samples (Figure 4C). To define the functional consequences of the *de novo* variants, high confidence variants were annotated using SnpEff (Figure 4D)(27). As expected, the majority of *de novo* variants occur within intergenic and intronic regions (Figure 4D). We hypothesize that if *de novo* variants are driven by the CRISPR/Cas9, then we would observe *de novo* variants occurring across all of the edited lines. We observed 19 *de novo* SNPs that occurred in all edited iPSC lines, none of which are predicted to be functional (Table 2; Supplemental Tables 1 and 3). These SNPs represent variants across the genome (Figure 2B). Given the minimal enrichment of shared, *de novo* variants across edited iPSC lines, we propose that off-target effects induced specifically by CRISPR/Cas9 are minimal.

**Figure 4:**
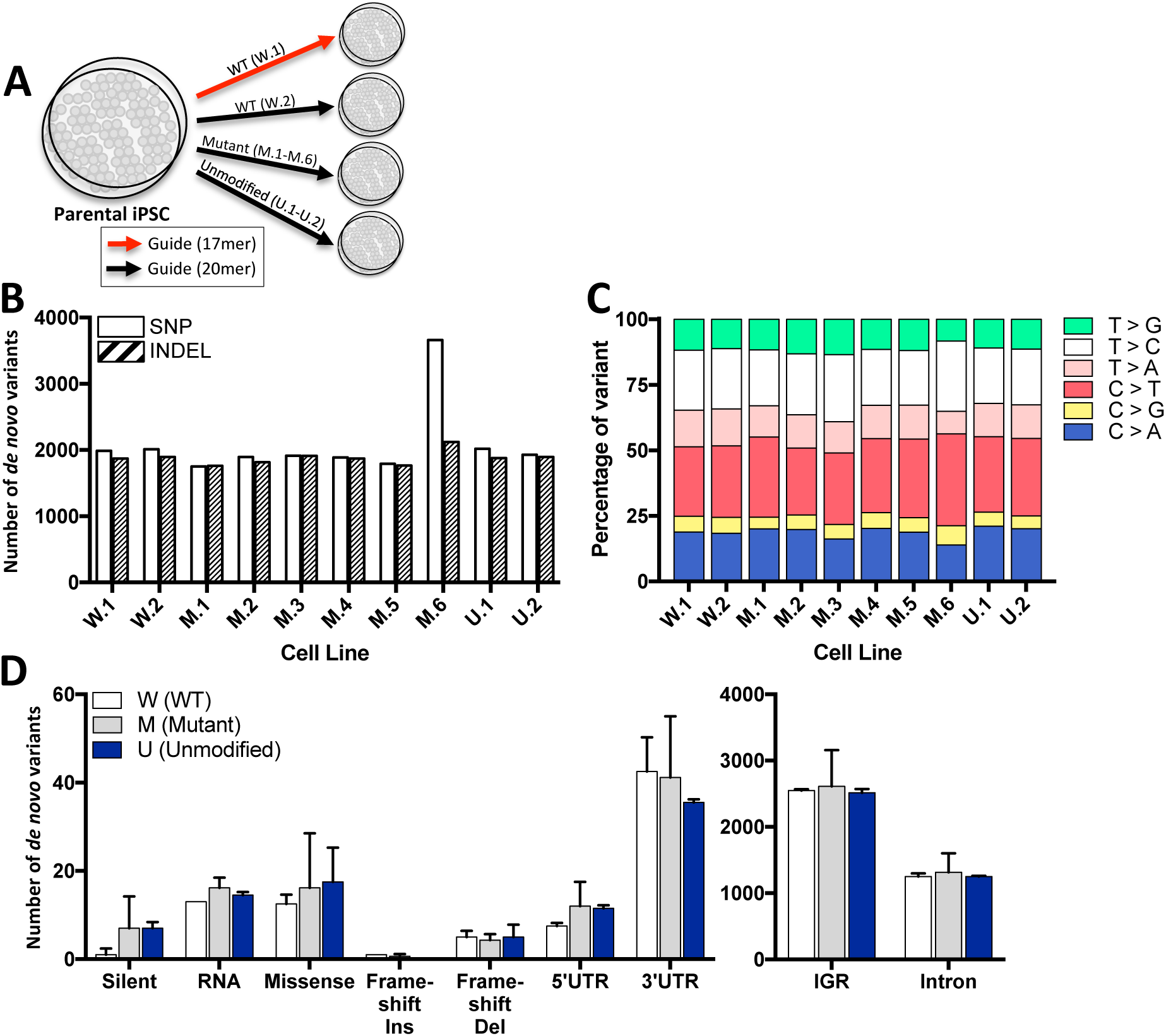
Summary of *de novo* variants in edited iPSC lines. A. Analyses included all *de novo* variants relative to parental iPSC (including those reported in dbSNP). B. Number of *de novo* variants: SNPs (white) and INDELs (hashed). C. Percentage of base changes reveal an absence of enrichment for a given base. D. Types of modifications within WT (white), mutant (gray) and unmodified (blue). W, mutant iPSC line modified to wild-type. M, mutant iPSC line modified to new mutation. U, mutant iPSC line unmodified.

**Table 2:** *De novo* variants present in exactly one or more edited iPSC lines

| **Variant Counts** | **Reference** | **SNPs** | **INDELs** | **Functional** | **Functional (%)** | **dbSNP** | **dbSNP (%)** |
| --- | --- | --- | --- | --- | --- | --- | --- |
| 1 edited line | Parental iPSC | 8,209 | 3,873 | 71 | 0.6 | 6,799 | 56.3 |
| 2 edited lines | Parental iPSC | 1,470 | 1,233 | 7 | 0.3 | 1,151 | 42.6 |
| 3 edited lines | Parental iPSC | 602 | 654 | 5 | 0.4 | 620 | 49.4 |
| 4 edited lines | Parental iPSC | 701 | 503 | 8 | 0.7 | 500 | 41.5 |
| 5 edited lines | Parental iPSC | 259 | 358 | 4 | 0.6 | 278 | 45.1 |
| 6 edited lines | Parental iPSC | 157 | 284 | 0 | 0.0 | 224 | 50.8 |
| 7 edited lines | Parental iPSC | 106 | 230 | 1 | 0.3 | 172 | 51.2 |
| 8 edited lines | Parental iPSC | 121 | 188 | 0 | 0.0 | 162 | 52.4 |
| 9 edited lines | Parental iPSC | 106 | 174 | 0 | 0.0 | 177 | 63.2 |
| 10 edited lines | Parental iPSC | 19 | 30 | 0 | 0.0 | 32 | 65.3 |

We propose that the majority of mutational burden observed in the edited iPSC is driven by *de novo* changes that occur when passaging cells. To test this hypothesis, we examined the mutational burden in iPSC that were exposed to the full genome-editing pipeline but which remained unmodified (reference: parental iPSC; Figure 3; Table 2). We observed 2,000 *de novo* SNPs in the unmodified iPSC clones relative to the parental iPSC lines, including between 10 and 20 of which are predicted to be functional (Figure 3; Supplemental Table 1). The majority of functional variants occur in genes that are known to produce false positives (*MUC, OR2, or FAM*). Pathway analysis reveals that the genes in which these variants occur are marginally enriched in pathways associated with immune function. This finding is consistent with the presence of *de novo* variants in genes in the HLA family where the size and recombination frequency makes WGS alignment and mapping challenging (Supplemental Table 2). Thus, we propose that selective pressures occurring during iPSC passage are driving mutational burden to a greater extent than CRISPR/Cas9-mediated off-target effects.

### Enrichment of de novo variants in dbSNPs

The majority of WGS analysis pipelines for iPSC include a step that removes all variants present in dbSNP prior to comparing parental and edited lines (6,17,23,24). However, variants reported in dbSNP represent rare Mendelian mutations, non-synonymous variants, and common variants that increase risk for complex diseases. Importantly, we found that *de novo* variants were significantly overrepresented at sites previously reported in dbSNP sites (Fisher’s exact test: p<2.2×10^−16^; OR 16.08; 95% CI 15.51-16.68; Figure 5; Supplemental Table 4). These findings indicate that genomic sites reported in dbSNP represent regions that are more susceptible to change. However, we cannot exclude the possibility that the current methods for calling and analyzing WGS may be weighted in favor of variants within dbSNP, resulting in enrichment of *de novo* mutations at these sites.

**Figure 5:**
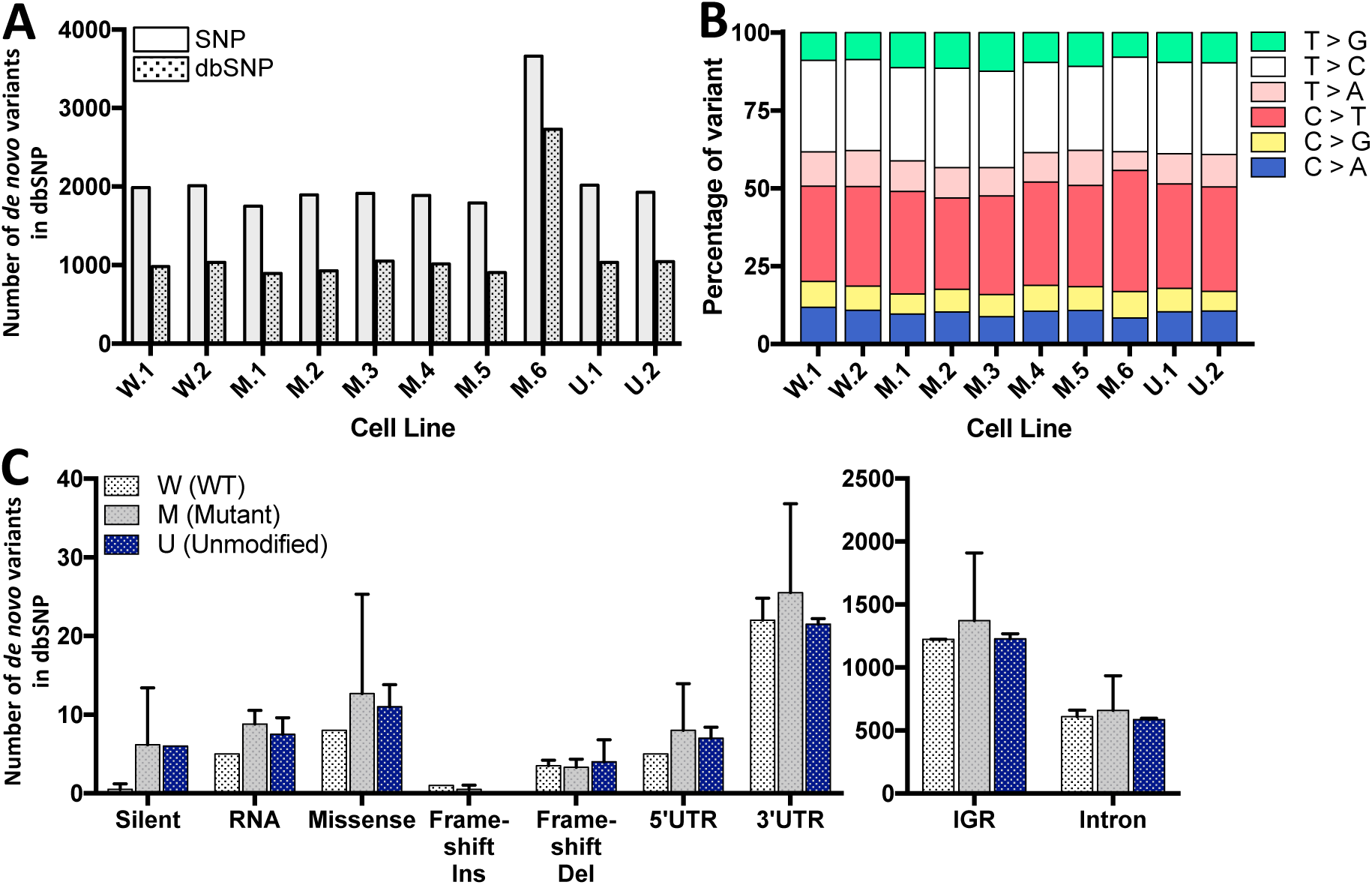
Enrichment of dbSNP among *de novo* variants identified in edited iPSC lines. Analyses included all *de novo* variants relative to parental iPSC that are reported in dbSNP. A. Number of *de novo* variants: SNPs and INDELs. B. Percentage of base. C. Types of modifications. W, mutant iPSC line modified to wild-type. M, mutant iPSC line modified to new mutation. U, mutant iPSC line unmodified.

## Discussion

Herein, we describe a highly efficient and robust method for allele-specific targeting by engineering allele-specific gRNA for CRISPR/Cas9 genome-editing in human iPSC. When designing gRNAs to modify mutant alleles in human iPSC for disease modeling, we have found that designing the gRNA sequence to overlap with the allele to be edited greatly improves our ability to target the mutant allele and to obtain correctly modified clones (Figure 1A). We observed similar efficiencies when applying our allele-specific gRNA design approach to multiple loci and donor iPSCs (Table 1). Previous studies have demonstrated that introduction of synonymous changes into a donor oligodeoxynucleotide may increase the recovery of clones with desired modifications while reducing NHEJ events at the target site (8,10). It is proposed that once homologous recombination occurs, the synonymous change prevents or blocks targeting of Cas9 to the edited allele. However, when attempting to attribute changes in cellular and molecular phenotypes to a single allele, the addition of synonymous changes near the site of interest complicates interpretation of the findings. Synonymous changes in the genome are known to cause disease and/or contribute to increase risk for many diseases (reviewed in (28)). Thus, we hypothesize that similar to the reports using synonymous changes in the donor oligonucleotides, designing allele-specific gRNA allows for increased specify for the target allele. This approach will accelerate the generation of isogenic controls to study contributions of pathogenic mutations and risk variants to molecular and cellular phenotypes in human iPSCs.

In thorough analysis of the off-target impact of CRISPR/Cas9, we demonstrate that CRISPR/Cas9 induces minimal mutational burden. Instead, we observe a larger impact on *de novo* changes occurring with culture. Of those *de novo* variants identified, we observed an enrichment of SNPs among those previously reported in dbSNP, which could have a greater chance of being functional.

## Methods

### Fibroblasts

Dermal fibroblasts were maintained in fibroblast growth media (Lonza) supplemented with penicillin/streptomycin.

### iPSC generation and characterization

Human fibroblasts were transduced with non-integrating Sendai virus carrying the four factors required for reprogramming: OCT3/4, SOX2, KLF4, and cMYC (29,30). Cells that show morphological evidence of reprogramming were selected by manual dissection and cultured under feeder-free conditions. Human iPSC lines were characterized using standard methods (Supplemental Figure 1) (29). Each line was analyzed for pluripotency markers (OCT4A, SOX2, NANOG, SSEA4, TRA1-60S, TRA1-81) by immunocytochemistry (ICC) and real-time PCR; for spontaneous differentiation into the three germ layers (GFAP, AFP, β3-tubulin, FOX2A, Desmin, SMA) by ICC and real-time PCR; and for chromosomal abnormalities by karyotyping.

### Maintenance and expansion of iPSC

Human iPSC were cultured in mTesR1 in feeder-free conditions on Matrigel-coated tissue culture-treated plates. For routine passaging and unless otherwise noted below, iPSC were dissociated with Accutase for 3 minutes. Dissociated cells were collected in PBS and centrifuged at 750 rpm for 3 minutes. After media was aspirated, a portion of the cells were plated on new Matrigel-coated plates in mTesR1 supplemented with 1uM Y-27632 (ROCK inhibitor). ROCK inhibitor was removed after 24 hours.

### Guide RNA design and validation

CRISPR reagents were obtained from the GEiC at Washington University School of Medicine (a gift from Keith Joung; Addgene plasmid 43860). Guide RNAs were designed to have at least 3bp of mismatch to any other gene in the human genome. Validation was performed using a mismatch detection assay to determine cutting efficiency (T7E1 assay) in K652 cells. The p3s-Cas9HC Cas9 expression plasmid was used (a gift from Jin-Soo Kim; Addgene 43945)(31).

### Single cell passaging of iPSC

Prior to nucleofection of the iPSC with the CRISPR/Cas9 reagents, human iPSC, which are typically passaged in small clusters, must be acclimated to culture as single cells. Because human iPSC cultured as single cells may exhibit increased cell death and/or spontaneous differentiation, we supplemented Matrigel with fibronectin and cultured the iPSC as single cells (via dissociation with Accutase for 10 minutes) for 2 passages. Fibronectin is a component of basement membranes and effective in maintaining rounded iPSC morphology upon passaging (32). Additionally, human iPSC derived from unique donors exhibit subtle variability in recovery from passaging as single cells. We have found that supplementing the Matrigel with fibronectin minimizes this variability.

To prepare cells for genome-editing, iPSC are serially passaged as single cells and maintained on Matrigel-coated plates supplemented with 5ug/cm^2^ fibronectin. To achieve single cell culture, iPSC were dissociated in Accutase for 10 minutes to achieve single cells and plated on Matrigel-coated plates supplemented with fibronectin in mTesR1 supplemented with 5uM ROCK inhibitor. Once the iPSC reach confluency, this process was repeated. This step is critical to allow the iPSC to acclimate to the single cell dissociation that is required for nucleofection and reduces spontaneous differentiation of the iPSC.

### Nucleofection

Upon reaching confluency, acclimated iPSC were nucleofected. We have found that two aspects are critical for successful nucleofection: (1) cell number and (2) relative concentrations of Cas9 and gRNA. After two sequential single cell passages, to prepare cells for nucleofection, iPSC were dissociated in Accutase for 10 minutes to achieve single cells and plated on Matrigel-coated plates supplemented with fibronectin in mTesR1 supplemented with 5uM ROCK inhibitor at a density that allowed for at least 5x10^6^ cells within 48 hours for nucleofection. Nucleofection was performed using a Lonza 4D-Nucleofector System. Briefly, iPSC were dissociated for 10 minutes in Accutase to achieve a single cell solution. The cell density was then measured using a Countess II Automated Cell Counter.

For the nucleofection reaction, 3x10^6^ cells were transferred to a 1.7mL tube. A control nucleofection reaction to assess sensitivity of the iPSC to the reaction used 1x10^6^ cells in a 1.7mL tube. The cell solutions were then centrifuged at 90xg for 5 minutes. Cells were then combined with the 1ug pMaxGFP (used to assess nucleofection efficiency), 1ug gRNA, 3ug Cas9, and 300uM single stranded donor oligo and the P3 Primary Cell 4D reaction mix (Lonza). For the control nucleofection reaction, cells were combined with 1ug pMaxGFP and the P3 Primary Cell 4D reaction mix. Cells were nucleofected using the Lonza program CA-137. After nucleofection, the cell solution was transferred to a one 35mm Matrigel-coated well that was pre-incubated with Recovery Media (DMEM/F12 supplemented with 10% FBS and 10uM ROCK inhibitor).

Nucleofection with fewer cells results in high toxicity and poor recovery 24 hours after plating. We tested several concentrations of Cas9 and gRNA. Nucleofection with higher concentrations of Cas9 and gRNA increased NHEJ events without proportionally increasing correctly modified clones. We have also observed differential sensitivities of independent iPSC donors to nucleofection; thus, the 3ug/1ug condition is sufficient to produce modified clones without excessive cytotoxicity in the more sensitive iPSC lines.

The recovery and proliferation period after nucleofection was found to be particularly critical for minimizing mosaicism and achieving clones with desired modifications in an efficient manner (Figure 1C and D). Nucleofected iPSC were maintained in the 35mm dish for 5-7 days. Within 24 hours of nucleofection, the FBS-containing media was replaced with mTesR1. During this time, the iPSC develop an amorphous morphology; however, transition to mTesR in subsequent days results in restoration of stereotypical iPSC morphology. Culturing cells in mTesR supplemented with ROCK inhibitor immediately after nucleofection resulted in extensive cell death, very little material for screening, and no successfully modified clones. Plating cells at lower density, for example 3 million cells in a 10cm dish, also increased cell death, suggesting that high cell density and serum supplemented media is critical for recovery of iPSC post-nucleofection. Over the 72 hours post-nucleofection, ROCK inhibitor was slowly reduced from the media. After 5-7 days, the iPSC were passaged for clonal isolation. This 5-7 day period represents a critical window in which Cas9 is active and allows for completion of Cas9-mediated cleavage prior to isolation of clonal populations and minimizes mosaicism.

### Serial dilution and clonal isolation of iPSC

To prepare cells for clonal isolation, nucleofected iPSC were dissociated for 10 minutes in Accutase to achieve single cells. Cells were then plated serially onto Matrigel-coated plates supplemented with fibronectin: approximately 30,000 cells are plate in the first well followed by a 1:10 serial dilution in two additional wells. The remaining iPSC that were not plated were frozen down in 2x iPSC Freezing Media (FBS supplemented with 20% DMSO) and mTesR1.

Approximately 7 days after serial dilution, clones were selected for picking. Ideal clones were ones that were large enough to survive the picking, physically isolated from neighboring clones, and producing homogenous pluripotent morphology. IPSC clones fitting these criteria were picked and transferred into a 96 well Matrigel-coated plate containing mTesR1 supplemented with 10uM ROCK inhibitor. Each clone was maintained independently in a single well.

Approximately 7 days after manual isolation of the iPSC clones, cells were sufficiently confluent within the wells to passage the cells for screening and storage. Ideally, the majority of wells will have iPSC that are between 40-80% confluent at the time of passage. To passage the iPSC, media from the individual wells are aspirated and the cells were washed with PBS. Next, 25ul of 0.05% trypsin-EDTA is added to each well and incubated at 37°C for 5 minutes. After the 5-minute incubation, 60ul mTesR is added to each well and the plate was tapped to mix. For DNA, 15ul of solution is transferred to a 96 well PCR plate while mirroring the location of each sample. For the remaining cell solution in the 96 well tissue culture plate, 50ul 2x iPSC Freezing Media is added and the plate was tapped to mix. To effectively store the plate for up to 4 months, the plate was wrapped in parafilm, lab tape was then wrapped around the parafilm, and the plate was stored in a Styrofoam box with any open areas filled with paper in the - 80°C. Securing the plate with both parafilm and tape is critical to maintain the seal and for enhancing the recovery of the cells upon thawing.

### Screening iPSC clones

A critical aspect of efficient editing is minimizing the reagents and effort required to maintain clones during the screening process. To screen iPSC for editing events, genomic DNA was extracted from IPSC in the 96 well PCR plate using QuickExtract DNA solution and following the manufacturer’s protocol. Genomic DNA was first assessed for the desired editing event by restriction fragment length polymorphism (RFLP). Briefly, primers were designed to amplify approximately 200 bp around the edited locus. The resulting PCR product was then incubated with a restriction enzyme that was identified to uniquely cut either the mutant allele or wild-type allele and the reaction was run on an agarose gel.

Clones that produced a restriction digest pattern consistent with the desired editing event were then selected for Sanger sequencing. At this time, we also selected 2-3 clones that appeared to be unmodified to serve as controls for exposure to the CRISPR/Cas9 and the editing pipeline. Briefly, 250bp on either side of the on-target editing event were amplified and sequenced using BigDye terminator, version 3.1, cycle sequencing kit and analyzed on an ABI 3500 genetic analyzer. Sequence analysis was performed using Geneious software (http://www.geneious.com, (33)).

Based on the Sanger sequencing results, clones were selected for expansion and characterization. Clones, named by their location in the 96 well plate, were expanded from the frozen plate into sufficient material to characterize and freeze down more cells. Briefly, the frozen plate was carefully removed from the Styrofoam box and placed into a 37°C tissue culture incubator for 15-20 minutes to thaw, while monitoring the plate to avoid warming the cells beyond the thaw. To dilute the freezing media, the cell material was transferred from the appropriate well into a 1.7mL tube. To the tube, mTesR and 10uM ROCK inhibitor was added to achieve a 1:5 dilution and final volume of approximately 500ul. The resulting cell solution was added to a Matrigel-coated 24 well plate. Daily media changes were performed and cells were expanded to achieve the necessary amount for characterization and cryopreservation.

### Characterization of edited and unmodified iPSC clones

IPSC clones with the desired on-target edits and those that remained unedited, were expanded in culture and material was collected for additional characterization in order to (1) identify on-target and off-target events and to (2) confirm that pluripotency was maintained. In order to identify on-target and off-target events, genomic DNA was extracted from cell pellets using the Gentra Puregene Cell Kit. Sanger sequencing was then performed as described above for the on-target regions and the predicted off-target regions. Off-target sequence reactions were performed for sites in the genome with 0, 1, and 2 mismatches with the guide sequence. IPSC clones were fully characterized as described above, including karyotyping.

### Whole genome sequencing

Whole genome sequencing was performed at the New York Genome Center using 3ug DNA. Illumina libraries were prepared using the TruSeqDNA PCR-free protocol, and 150 bp paired-end reads were sequenced on the Illumina HiSeqX to an average depth of 30x autosomal coverage.

### Alignment of sequencing reads

Output reads were aligned to the NCBI GRCh37 reference genome using the Burrows-Wheeler algorithm (BWA; http://bio-bwa.sourceforge.net/) (34). Duplicate reads were marked by Picard (https://broadinstitute.github.io/picard/). Base quality score recalibration, local realignment around indels, and downstream variant calling were performed using the Genome Analysis Toolkit (GATK) v3.4-46 according to GATK Best Practices (https://software.broadinstitute.org/gatk/)(22).

### SNP and Indel calling

Multiple filtering steps were undertaken to produce a high quality set of variants. Variants were removed which did not pass variant quality score recalibration (GATK) (35). Variants were also removed at sites where the depth of coverage was excessively high (depth > mean + 3 standard deviations), and in low complexity regions (36). Genotype calls with low depth (DP < 8) or poor genotype quality (GQ < 20) were removed (37). Additionally, variants were removed where all CRISPR-Cas9 edited samples with called genotypes matched the reference fibroblast or iPSC line. Finally, complex variants (e.g. overlapping INDELs) were removed using GATK (Supplemental Figure 2). One edited line was dropped during analysis due to a high number of *de novo* variants occurring in chr18:32470291-77905340, suggestive of a chromosomal rearrangement in this region.

### Off-target analyses

Sites with a high potential of off-target editing by the CRISPR-Cas9 system were predicted (15). All variants within 300bp of these sites were extracted (GATK) and manually evaluated.

### Annotation

High quality variants were annotated using snpEff (27) using database GRCh37.75, dbSNP build 147 (http://www.ncbi.nlm.nih.gov/SNP/), the Exome Variant Server (http://evs.gs.washington.edu/EVS), and the 1000 Genomes Project (38).

Subsets of variants extracted using SnpSift (39), making extensive use of GNU Parallel to perform multiple operations simultaneously.

### Pathway Analysis

The primary gene associated with each variant was extracted using SnpSift, and a list of gene was analyzed using the *ToppGene* portal (https://toppgene.cchmc.org/).

### Identification of de novo variants in iPSC exposed to CRISPR/Cas9

Each edited line was compared in a pairwise genotype manner to the parental iPSC line. Counts were generated for nucleotide base change, variant class (coding, intron, etc.), and dbSNP membership for each variant using bcftools (40), SnpSift, and GATK, using annotated VCF (41) files.

### Plot of variant locations

The positions of variants in all edited lines were aggregated and plotted using R package ggplot2 (42).

## Author Contributions

JPB, RM and CMK designed the study and wrote the manuscript. RM and SH performed genome-editing. AMG provided conceptual contributions. JAC and GC generated WGS data. JPB and CC developed WGS analysis pipeline and analyzed data. NW performed pathway analysis.

## Acknowledgements

We would like to thank the Washington University Genome Engineering and iPSC Center for advice in the generation of guide RNAs. This work was supported by access to equipment made possible by the Hope Center for Neurological Disorders, and the Departments of Neurology and Psychiatry at Washington University School of Medicine. Funding provided by NIH-K01 AG046374 (CMK), Hope Center (CMK), Rainwater Charitable Foundation (CMK, AMG, GC). Fibroblast lines were a gift of the NINDS Cell Repository (F0510 and ND32951A).

